# Bleaching correction for DNA-measurements in highly diluted solutions using confocal microscopy

**DOI:** 10.1101/2020.04.06.027375

**Authors:** Lorenz T. Sparrenberg, Benjamin Greiner, Harald P. Mathis

**Affiliations:** Chair of Biotechnology, RWTH Aachen University, Aachen, Germany; Fraunhofer Institute of Applied Information Technology FIT, Sankt Augustin, Germany

## Abstract

Precise and reliable determination of the nucleic acid concentration in biological samples still remains a challenge. This is particularly since the established fluorescene-based methods provide insufficient results, when only minute sample quantitites are available for analysis. Among other effects, photobleaching is the main reason for this. Since large molecules diffuse more slowly than small molecules, they are exposed to more excitation cycles and therefore have a higher probability of permanently losing their fluorescence. Solutions with large molecules hence show a reduced fluorescence. In this paper we present a method to correct this effect and thus allow high-precision sample concentration determination in minute sample quantities (< 2 µl drops with concentrations < 20 pg/µl). For this purpose, we used confocal microscopy with single molecule sensitivity. In the first step, we derived calibration curves from DNA solutions with defined fragment length. We analyzed dilution series over a wide concentration range (1 pg/µl – 1000 pg/µl) and measured their specific diffusion coefficients by employing fluorescence correlation spectroscopy. Using this information, we correct the measured fluorescence intensity of the calibration solutions for photobleaching effects. Subsequently, we evaluated our method by analyzing a series of DNA mixtures of varying composition.

## Introduction

In molecular biology, precise knowledge of molecule concentrations, especially nucleic acid concentration, is required. Various molecular biological methods such as molecular cloning or sequencing involve nucleic acids and depend on precise concentration information [1,2]. Also, the analysis of tissue samples or the examination of expression patterns of cells require exact concentration information on the extracted nucleic acid amounts [3–5]. PCR based methods can analyze nucleic acid samples consisting of only a few templates and are therefore widely used in molecular biology [5,6]. However, these methods rely on sequence information from the sample. PCR based methods cannot aid in the analysis of nucleic acid mixtures of unknown sequences. Fluorescence measurements are highly promising for this task because of their extraordinary sensitivity that even allows measurements of single molecule events [7]. A number of fluorescent dyes is available that bind sequence independently to nucleic acids and thus enable reliable labelling [8,9]. Additionally, fluorescent dyes for labeling are inexpensive and easy to handle. To save as much of the valuable nucleic acid sample as possible, measurements in highly diluted solutions with microliter volumes are preferred [2]. Confocal fluorescence microscopy can meet these challenges and is therefore the ideal candidate for measuring mixtures of unknown nucleic acid sequences.

To obtain reliable results in fluorescence-based concentration determination, we have to consider various effects. In confocal measurements soaring power intensities occur which may be of the order of several 100 kW/cm^2^ [10], which of course exposes the fluorophores to an enormous load. This is why the phenomenon of photobleaching, whereby molecules can permanently lose their fluorescence property due to irradiation with excitation light [11], is particularly noteworthy. During confocal measurements on freely diffusing molecules, a stationary equilibrium between the particle streams of bleached and fluorescent molecules is established in the excitation volume, so that on average an apparently lower intensity, and thus also a lower concentration, of fluorophores in the solution is measured due to the bleaching effect.

The extent to which a measurement is influenced by photobleaching depends on various factors, such as the irradiance of the excitation light, the photochemical properties of the fluorophore used or the proportion of oxygen in the solvent [11]. The probability that a fluorophore is bleached is directly related to the duration of irradiation by the excitation light [12]. As a result, the bleaching rate occurring in the solution is a function of the molecular size, since large molecules diffuse more slowly than small molecules and thus remain longer in the excitation volume. The similar effect is valid for mixtures of nucleic acids with arbitrary fragment length distribution. The only difference is that the mean fragment length and the mean diffusion constant are the characteristic properties of the mixture for photobleaching. Consequently, nucleic acid mixtures of the same mass concentration but with different molarity have different mean diffusion constants and due to photobleaching have different fluorescence intensities.

We have derived a new calibration procedure to account for this effect and to correct the effect of photobleaching in fluorescence measurements. For this purpose, we use fluorescence correlation spectroscopy in a confocal setup to analyze the diffusion properties of DNA solutions. With this information, we can correct the fluorescence measurements for photobleaching to achieve highly accurate results.

Fluorescence correlation spectroscopy (FCS) is a commonly used technique to analyze the diffusion behavior of particles. The idea behind FCS is the correlation in time of thermodynamic concentration fluctuations of diffusing particles in solution in equilibrium [13]. Eq 1 gives the autocorrelation function to calculate the FCS and corresponds to the correlation of a time series with itself shifted by time *τ*, as a function of *τ*:

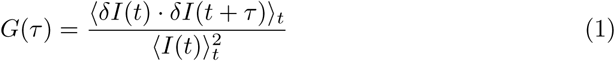

where *I*(*t*) is the measured intensity at time *t*. ⟨*…* ⟩_*t*_ denotes a time average over time:

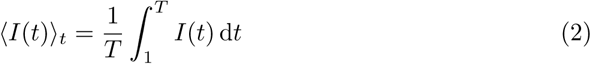

The intensity fluctuation *δI*(*t*) at a certain time is:

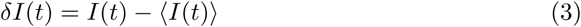

From this, the autocorrelation function turns into

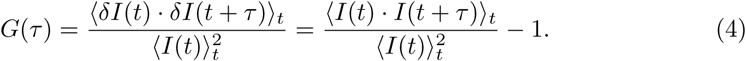

In this form, we can easily compute the autocorrelation from a given fluorescence trace.

On basis of physical considerations, a model for the autocorrelation can be derived. For this, the properties of the laser profile and molecular diffusion properties of the sample are considered. This model gives ground to approximate the detection efficiency of a diffusing particle excited by a single-mode laser in a confocal setup by a Gaussian profile. [15]:

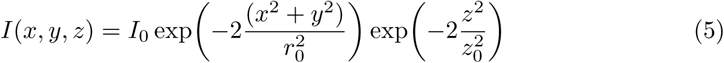

where *x, y* and *z* are the coordinates of the observation volume. *z* is the direction of the laser beam. *r*_0_ is the radius of the observation volume and (2 *z*_0_) is the effective length of the volume. For the Gaussian distribution of the laser profile, we can write the autocorrelation function for one freely diffusing particle species as follows [16]:

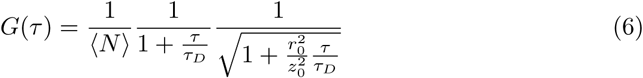

Here, *N* is the average molecule number in the detection volume. *τ*_*D*_ is the average diffusion time of particles in the observation volume giving the characteristic decay scale of fluorescence fluctuations [14]. We use equation 6 to fit the experimental FCS data to get the diffusion time *τ*_*D*_ from the measurement. At *τ* = 0, we obtain the mean number of diffusing particles in the volume ⟨*N*⟩ or equivalently the mean concentration ⟨*C*⟩:

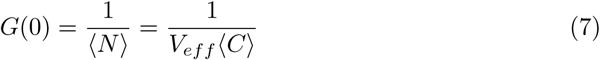

Where the effective observation volume is:

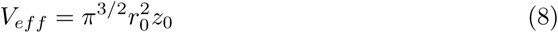

The diffusion coefficient of the diffusing molecules in solution is:

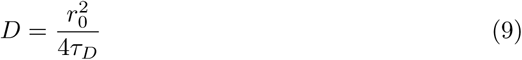

Standard implementations of FCS methods do not provide absolute values of diffusion coefficients, since they require information about the geometric shape of the detection volume, which is challenging to measure independently. Therefore, measurements of the diffusion coefficient are relative and require additional measurements of a reference substance e.g. Alexa 488 with known diffusion coefficient and concentration to derive the information about the geometric shape [14].

It is important to note that with small shifting times further effects like e.g. triplet state effects (microseconds) [17] or photodiode afterpulsing (nano-to microseconds) [18] dominate the autocorrelation and Eq 6 needs to be adapted. Since in the present work polymeric molecules with comparatively large diffusion times (*τ*_*D*_ *>* 1 × 10^−3^ s) were analyzed, the important changes of the autocorrelation function take place at relatively large shifting times and these effects acan be neglected.

## Materials and methods

For our experiments we used a home-build fluorescence microscope in a confocal setup (Laser: Lasos LDM F series, 90 mW, 486 nm; Filters: Linus 1% neutral filter, Bright Line^®^ fluorescence filter 535/40; Dichroid mirror: Linus 500 LP; Objective: Zeiss LD Plan-Neofluar 63x / 0.75 korr, ∞ / 0-1.5; Detector: Avalanche photo diode from Micro Photon Devices PDM series 100 µm. The pinhole has a diameter of 100 µm. An ALV correlator card processed the fluorescence signal. Fig 1 shows a schematic description of the confocal setup and the simplified measurement principle. For each fluorescence measurement, we placed a 2 µl drop on a cover slip and positioned it under the objective lens with the drop pointing away from the lens. Then we approached the measuring position along the optical axis, which was 50 µm in the drop volume. A pause of 30 s after each approach guarantied an almost reached equilibrium of bleached and unbleached fluorophores in the detection volume. The actual measurement took 30 s. During this time the ALV correlator card collected the data and calculates the autocorralation of the measurement. The data were fitted to equation 6 to get the characteristic diffusion times *τ*_*D*_ of each measurement. For the estimation of the geometric shape of the detection volume, measurements were conducted with the fluorescence dye Alexa 488 (PicoQuant) which has a known diffusion coefficient of 435 µm^2^s^−1^ [19].

**Fig 1.**
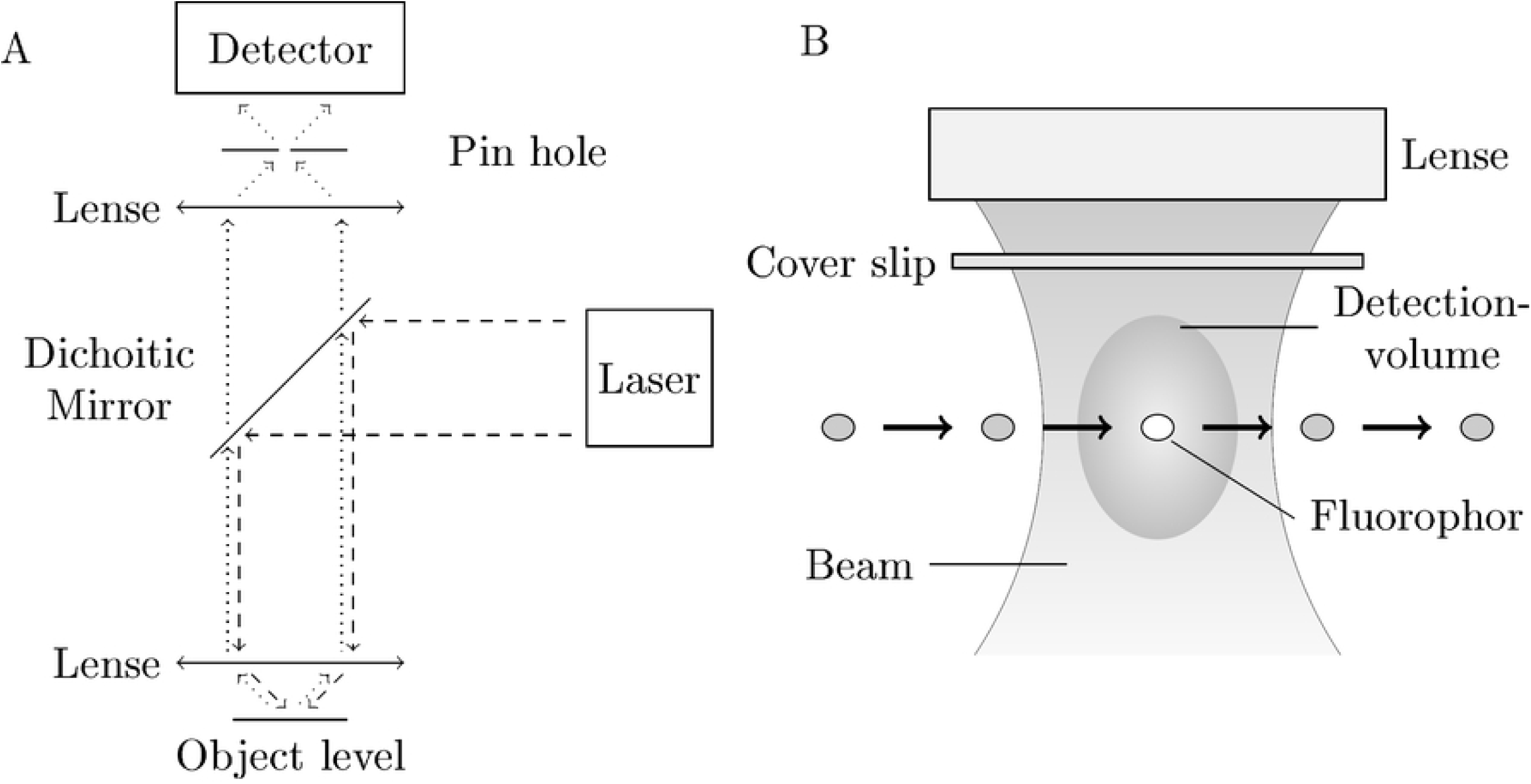
Schematic of the experimental setup. A: A laser emits light that is passed through a dichroic mirror and focused on the sample through an objective lens (dashed). Excited fluorophors in the probe start to emit photons which are collected by the objective lense. Because of their extended wave length these photons can pass the dichroid mirror. A detector counts these emitted photons (dotted). The preceding pinhole improves the signal to noise ratio and assures that only photon events from the object level can reach the detector. B: A fluorophor enters by random walk the detection volume and begins to emit photons. The average time to pass the volume correlates with its diffusion coefficient

For the calibration procedure, dsDNA solutions (NoLimits™, ThermoFisher Scientific) with fragment lengths of 50 bp, 200 bp, 500 bp, 1000 bp, 2000 bp, 3000 bp, 6000 bp and 10 000 bp in water/DMSO solutions with a mass fraction of 3 parts water in 1 part DMSO were prepared. Dilution series from 1 pg/µl to 1000 pg/µl for each fragment length were recorded whereas each measurement was conducted five times. During all measurements the temperature was constant at 22 °C.

For verification of our calibration procedure, we set up DNA mixtures of known composition. The first set of mixtures consisted of 200 bp and 500 bp dsDNA with the ratios 1:1, 1:2, 1:3, 1:4 and 1:5. The second set consisted of 50 bp and 1000 bp DNA with the ratios 1:1, 1:2, 1:3, 1:4, 1:5 and 1:6. For each ratio, we prepared concentrations of 20 pg/µl, 50 pg/µl, 100 pg/µl and 200 pg/µl in water/DMSO solutions with a mass fraction of 3 parts water in 1 part DMSO. Again, we performed each measurement five times.

## Results and discussion

To derive a calibration procedure to correct photobleaching effects, we started with analyzing the diffusion properties of DNA-solutions. Therfore, we used fluorescence correlation spectroscopy (FCS) to gather the required information. Fig 2 shows the results of the FCS measurements exemplarily for the 50 bp dilution series. The black solid lines correspond to the fit of Eq 6 to the measurement data to get the specific diffusion time *τ*_*D*_ for each fragment length. We have only taken into account the data for shifting times greater than 1 × 10^−4^ s in order to bypass additional effects such as triplet state effects or APD afterpulsing which occur at small shifting times of nano-to microseconds. For increasing concentration the amplitutes of the graphs become smaller, whereas *τ*_*D*_ remains constant for all graphs. The FCS results for the other fragment sizes are comparable to the shown results with its corresponding diffusion times *τ*_*D*_.

**Fig 2.**
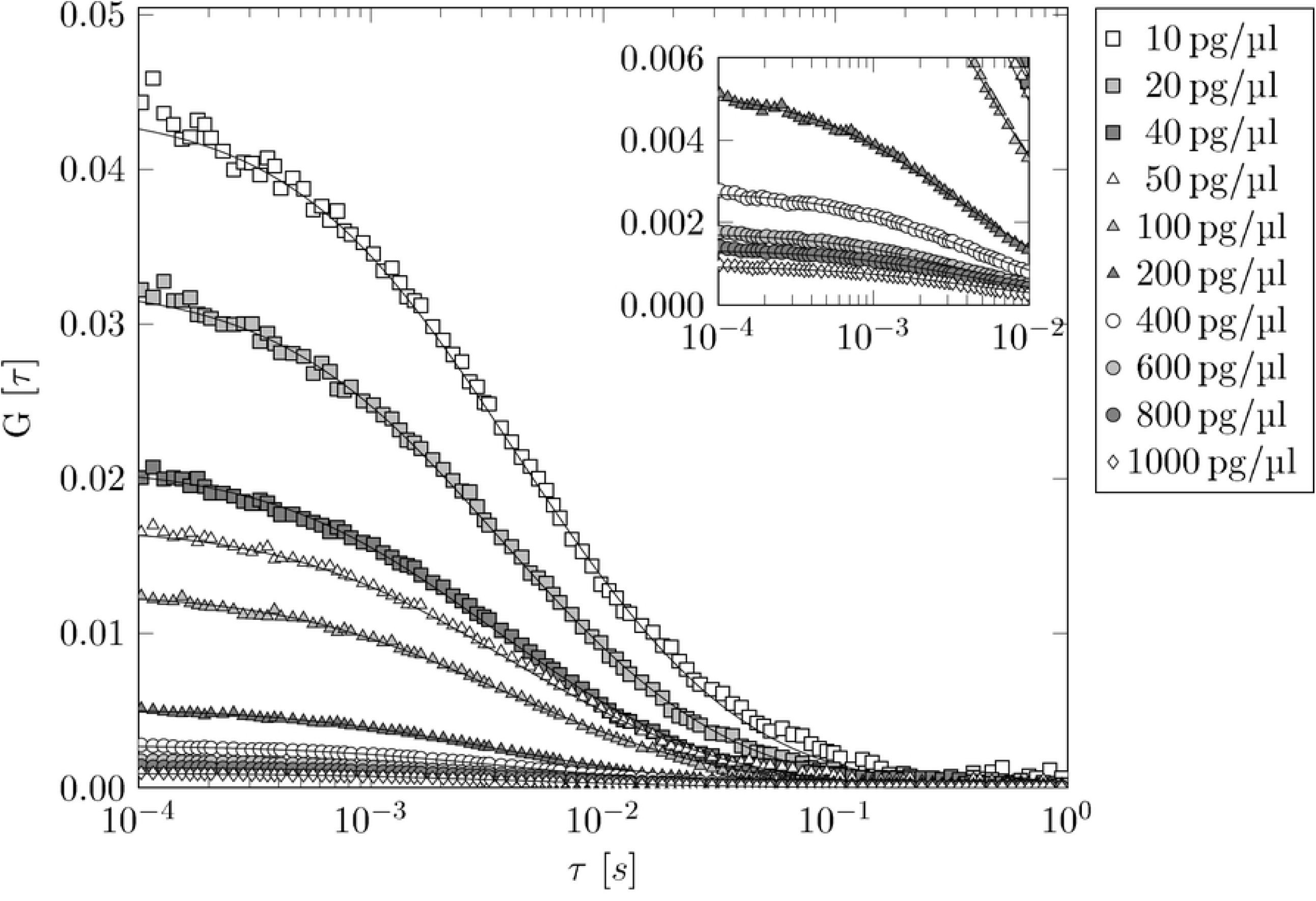
FCS data of 50 bp DNA dilution series. For clearity, we only show the median of the five single measurements of each dilution step. The black solid lines correspond to the fit of Eq 6 to the data in order to get the specific *τ*_*D*_ values for each fragment length. For 1 pg/µl DNA-solution (not shown) the FCS yields implausible results because of to less fluorescence events in the detection volume.

With the geometric information about the excitation volume of the laser we can now calculate the diffusion constants for each fragment size using equation 9. To obtain the geometric information of the observation volume we conducted FCS measurements with solutions of known diffusion coefficients. In our case, we used Alexa 488 for this purpose. According to Zimm’s model for flexible polymers in solution with excluded volume assumptions, we expect for DNA-solutions an exponent of around −0.60 because of the scaling law *D ∝ M* ^−0.60^ [20] where *D* is the diffusion coefficient and *M* is the molecular weigth. Actually, we found an exponent of −0.567 which is very close to previously reported values of −0.57 [21] and −0.571 ± 0.014 [22] for dsDNA molecules in aqueous solution (see figure 3). The exponent from the scaling law is independent of the ambient temperature during the experiments. Therefore, we can directly compare the exponents.

**Fig 3.**
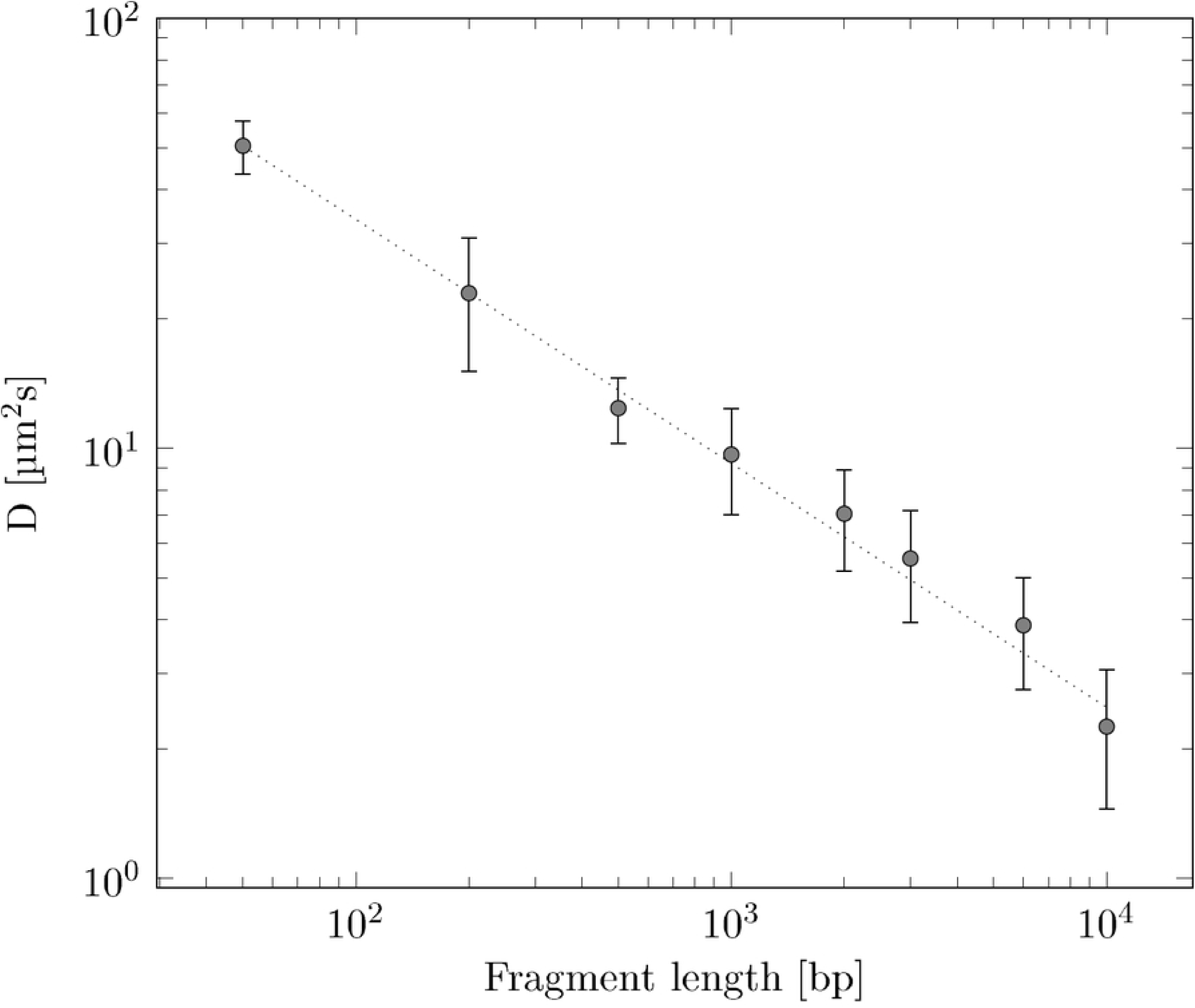
Diffusion coefficients of DNA solutions of different fragment sizes. The data were taken at 22 °C. The square marks correspond to the measurement data whereas the dotted line corresponds to the fitted model (*f* (*bp*) = *a* × *bp*^*b*^ with *a* = 494.065 and *b* = −0.567

The results of the diffusion measurements and their comparison to literature prove the reliability of our measurements. We would like to point out that we have used the diffusion time *τ*_*D*_ instead of the diffusion coefficient *D* in the following sections.

The next step in our method involves the calculation the median of the fluorescence intensity over time for each fragment length and each dilution step. Fig 4 shows exemplarily the results for 50 bp DNA.

**Fig 4.**
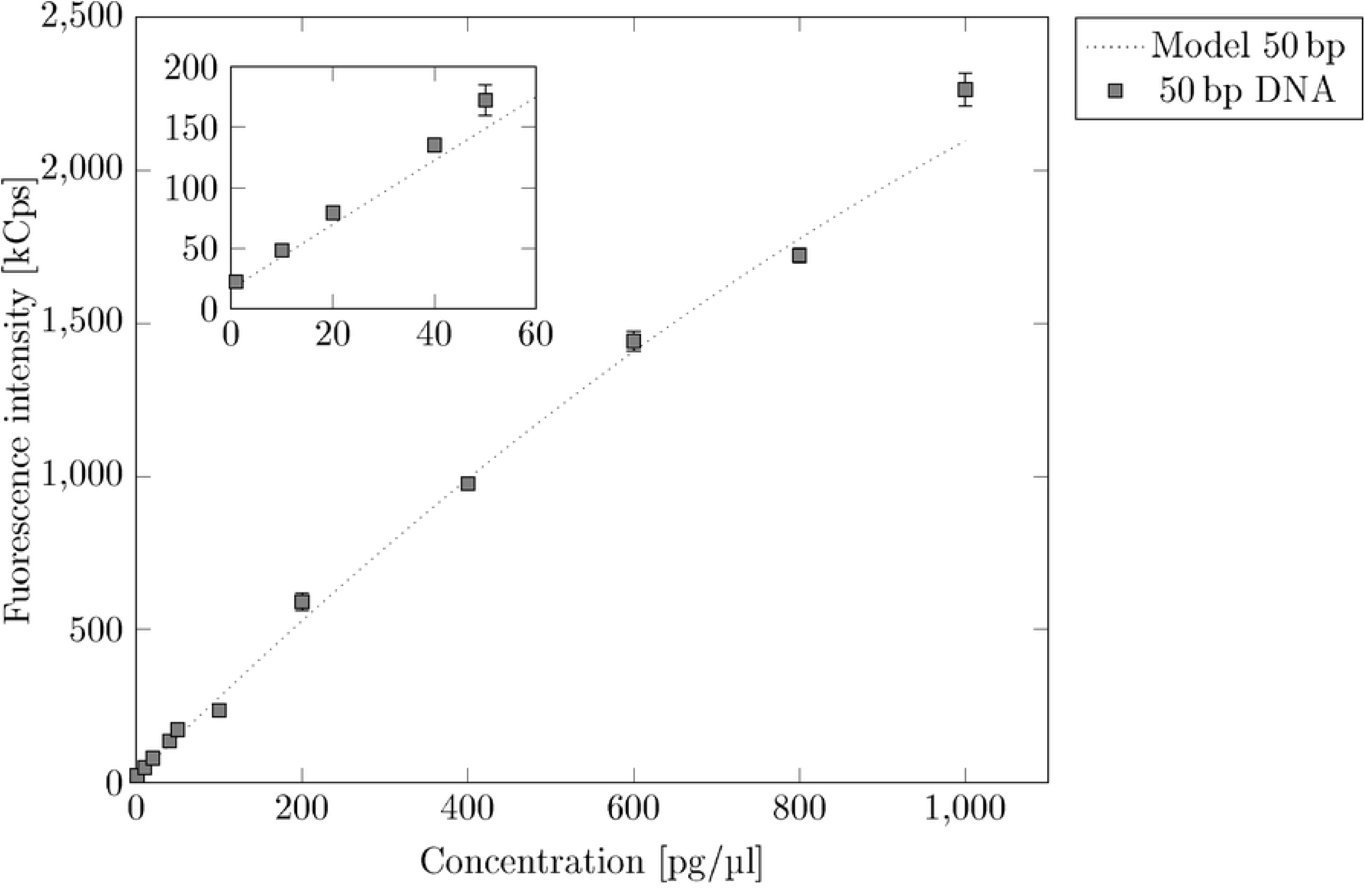
Intensities of 50 bp DNA dilution series. We used a polynome with the shape *I* = *f* (*C*) = *aC* + *bC*^2^ + *const* to fit the data. With *const* = 16.417, we get *a* = *a*_*cal*_ = 2.68 and *b* = −0.0006.

The resulting graph is almost linear and we can fit the data with a straight line with sufficient accuracy. At high concentrations, however, the graph is best described by a polynomial of 2nd order, since the quadratic term takes into account concentration-dependent effects such as quenching or volume exclusion. Therefore, we fit the dilution series of 50 bp DNA to a polynomial function the form

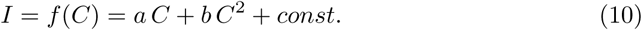

*I* is the intensity and *C* is the concentration of the analysed solution. The *y*-axis intersections *const* is set to the backgound noise we observed during our measurements (16.417 kCps). Next, we rotate the fitted function *I* = *f* (*C*) around the *z*-axis intersection *const* to map the data of the other DNA-solutions of different fragment size. For this purpose we multiply the vector 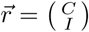 with a rotation matrix:

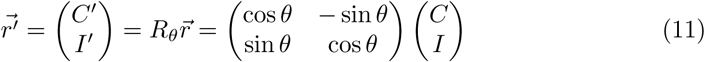

where *C*′ and *I*′ are the measured concentration and intensity affected by photobleaching. By inserting *I* = *f* (*C*) in Eq 11 and by translating it into the origin,

we obtain for the expressions *C*′ and *I*′:

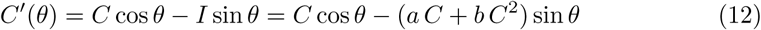

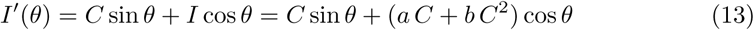

Now, we want to express Eq 13 as a function of *C*′. Therefore, we need to resolve Eq 12 to *C*:

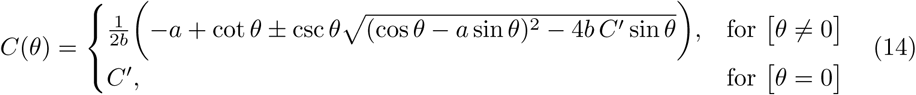

In our case *θ* lies between 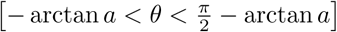 because physically only values in the first quadrant (positive concentrations and positive fluorescence rate) are reasonable. Furthermore, we only consider positive values for *a* and concentration *C* while *b* has to be minimal and negativ (|*b*| << *a*). Last but not least, only the negativ term of Eq 14 is reasonable. Thus, we discart the positive term of Eq 14. Now, by inserting Eq 14 into Eq 13 and by translating the expression back to *const*, we get the following expression:

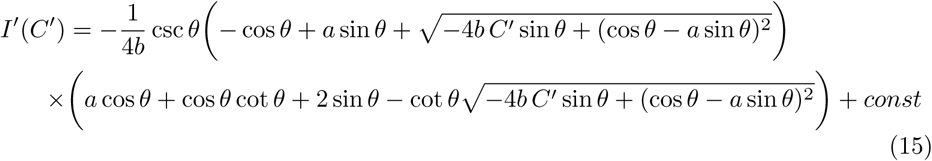

Eq 15 rotates Eq 10 around the *z*-axis intersection *const*. The validity of the approach is limited to functional areas where the rotated Eq 10 is monotonously growing. For larger concentrations the quadratic term starts to dominate and the approach is no longer valid. Eq 15 is subsequently fitted to the data of the remaining DNA dilution series (200 bp, 500 bp, 1000 bp, 2000 bp, 3000 bp, 6000 bp, 10 000 bp) to get the rotation angle *θ* for each fragment length (see Fig 5). We are aware of the fact that we can fit each dilution series directly to a polynomial function without the detour via rotation. But the procedure using one calibration curve and rotating it to fit the data seems to be much more stable. The resulting rotation angles *θ* for each dilution series provides the corresponding slope *a*_*sample*_. For this we rotate slope *a* around the angle *θ*.

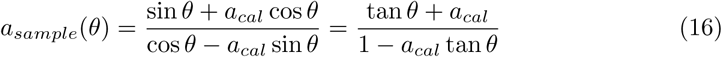

**Fig 5.**
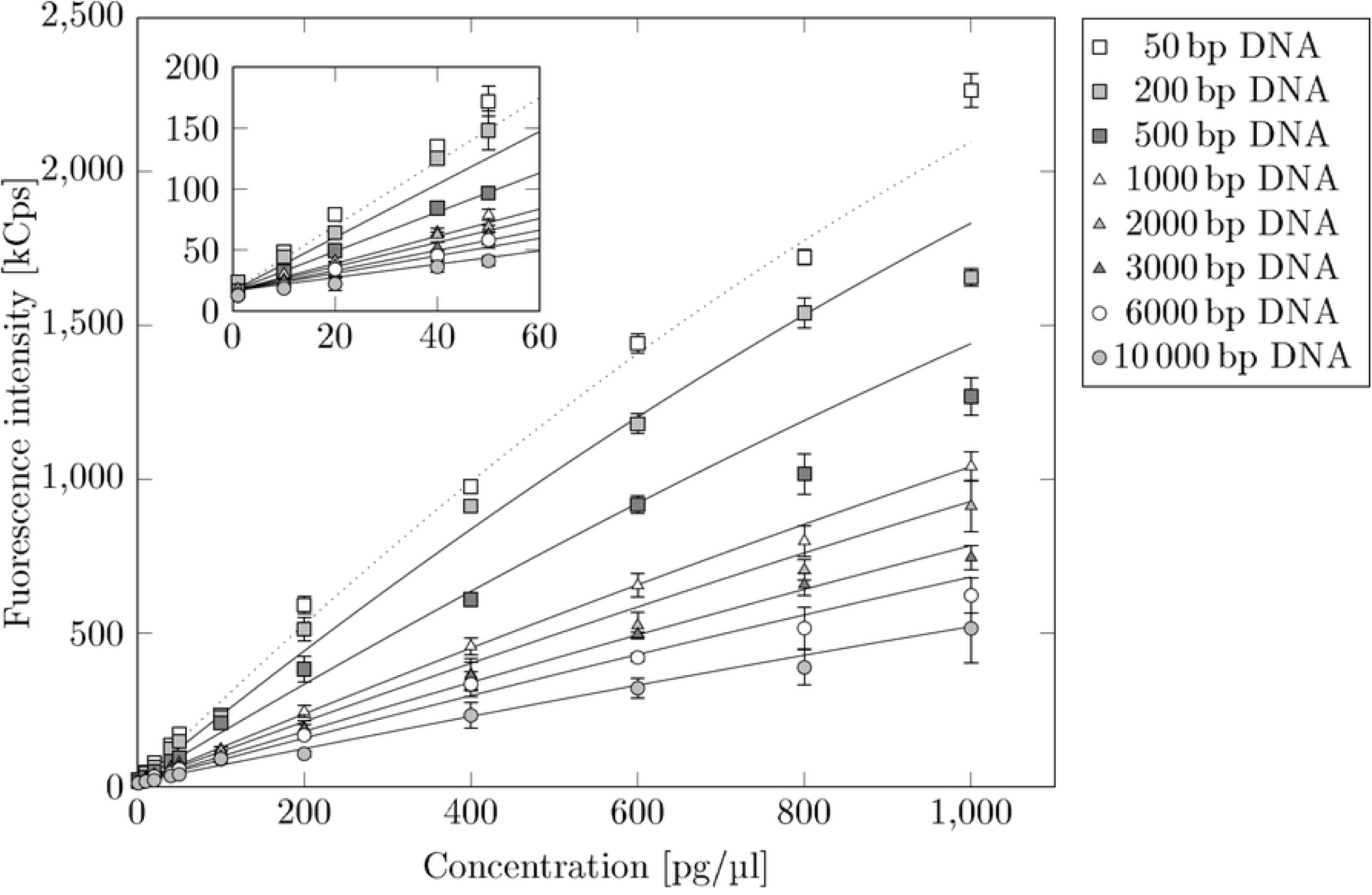
Bleaching effect on dilution series of different DNA fragment sizes. Dilution series of 50 bp, 200 bp, 500 bp, 1000 bp, 2000 bp, 3000 bp, 6000 bp and 10 000 bp DNA fragments. We rotate the polynomial fit (dotted line) of 50 bp DNA clockwise to map the other dilution series.

Here, the slope *a*_*cal*_ = *a* comes from the 50 bp dilution series in Fig 4. Now, we plot the resulting slope of each rotated curve against its diffusion time, respectively. The result can be seen in Fig 6. For modelling of the photobleaching as a function of *τ*_*D*_ we employed a propability-based approach:

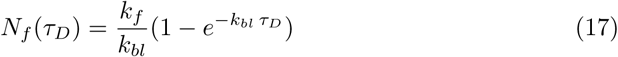

**Fig 6.**
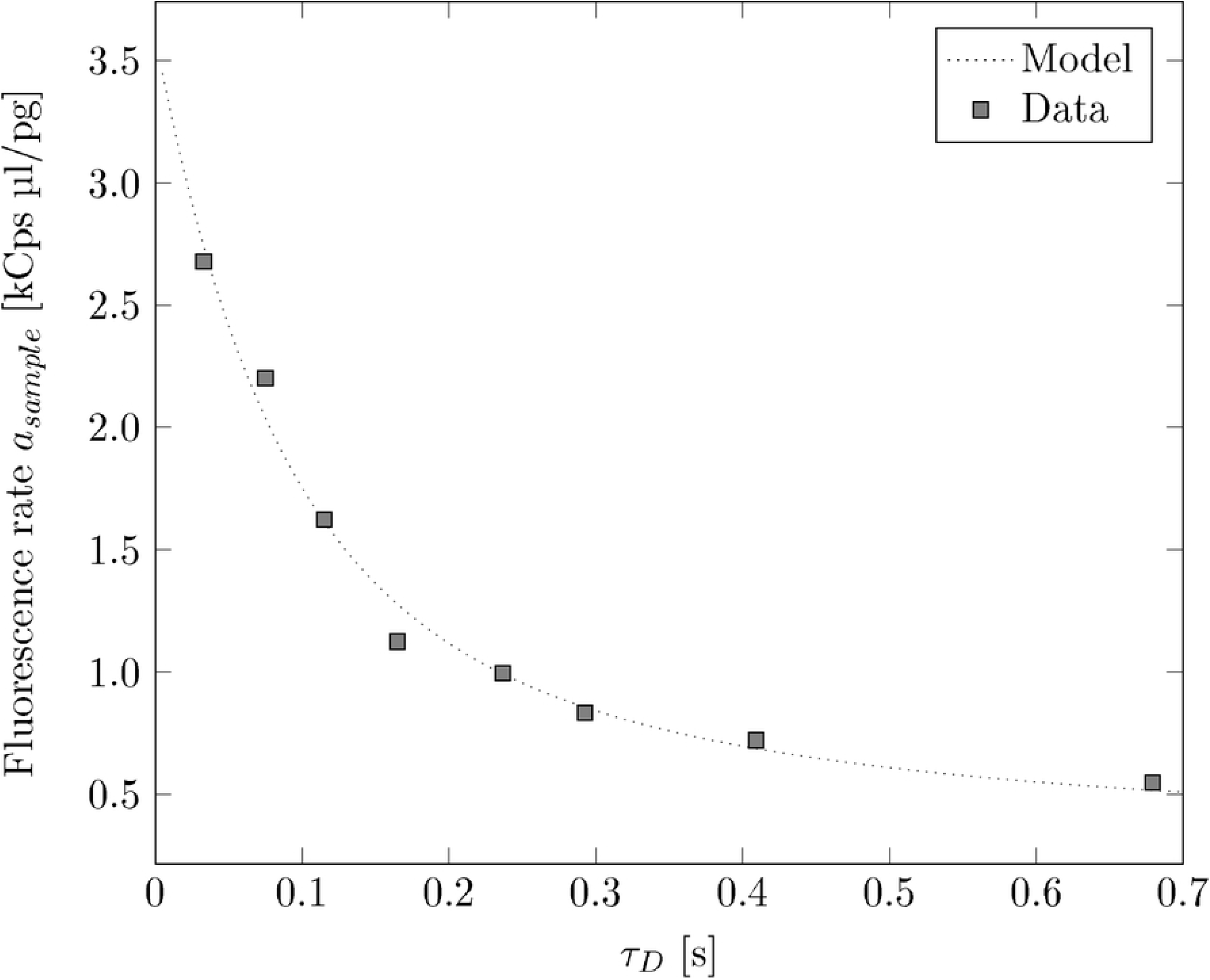
Slopes of the bleached dilution series against the diffusion times. The fluorescence rate as function of diffusion time for 50 bp, 200 bp, 500 bp, 1000 bp, 2000 bp, 3000 bp, 6000 bp and 10 000 bp DNA fragments. The fit Eq 18 to the data yields *k*_*int*_ = 0.176, *k*_*bl*_ = 19.016 and *const* = 0.257. The photobleaching effect affects the fluorescence rate which is hence lower for bigger diffusion times (larger molecules).

Here *N*_*f*_ is the average number of fluorescence photons and *k*_*f*_ and *k*_*bl*_ are the fluorescence rate and bleaching rate, respectively [23]. By setting 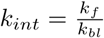 and dividing the expression by *τ*_*D*_, we obtain the rate of fluorescent photons depending on the diffusion time. Finally, the introduction of a constant *const* is necessary to take into account the fact that the fluorescence rate for long diffusion times can never become zero but is approaching a limit value. Putting all these considerations together, Eq 17 turns into:

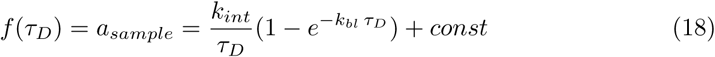

The fit of Eq 18 to the data yields: *k*_*int*_ = 0.1759, *k*_*bl*_ = 19.0164 and *const* = 0.2571 (see Fig 6).

## Determination of concentration and evaluation of method

In order to determine the exact mass concentration of a DNA sample of unknown composition, the following steps are performed:

1. Measuring of fluorescence intensity *I*′
2. Determination of *τ*_*D*_ via calculation of the autocorrelation
3. Calculation of *a*_*sample*_ using Eq 18
4. Determination of the angle of rotation *θ* via *a*_*sample*_ and *a*_*cal*_ from the calibration (50 bp DNA) using Eq 16
5. Calculation of the corrected concentration *C* via Eq 15 and *I*′

The fluorescence intensity *I*′ is obtained directly from the measurements. The mean diffusion time *τ*_*D*_ for a sample is then determined from the measurements using the autocorrelation. Eq 18 gives the characteristic slope *a*_*sample*_ for a given diffusion time *τ*_*D*_. Now, resolving Eq 16 to the angle of rotation *θ* yields the reverse function *θ*:

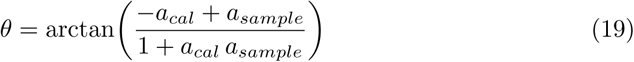

By inserting *a*_*sample*_ from the step before and *a*_*cal*_ from the 50 bp calibration measurement, we get the resulting rotation angle *θ* of the sample. To calculate the actual concentration of the DNA sample, we use the reverse function *C*(*θ*) of Eq 15:

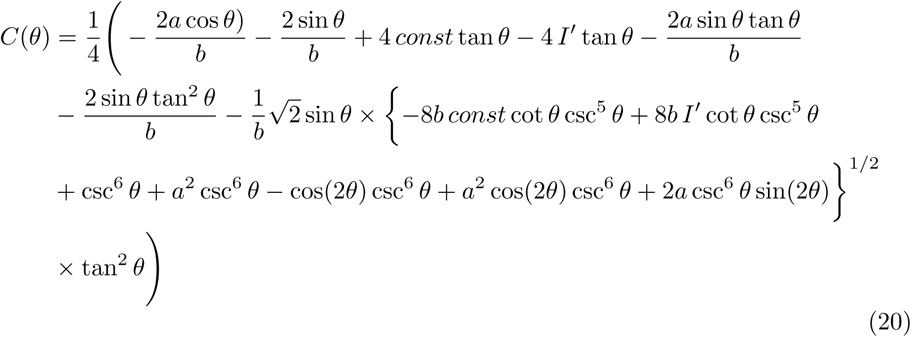

Inserting the calculated angle *θ* and the measured intensity *I*′ of the sample yields the actual mass concentration of the DNA sample.

For evaluation of the procedure, we used a set of 11 DNA mixtures with the concentrations 20 pg/µl, 50 pg/µl, 100 pg/µl and 200 pg/µl, respectively. We conducted each measurement five times and analyzed the fluorescence traces by means of FCS to get the corresponding diffusion time *τ*_*D*_. We averaged the results of the fivefold measurements by using the median and calculated the corrected concentrations for each DNA mixture using the above schematic.

The deviation from the setpoint concentrations are below 2.3% for 50 pg/µl, 100 pg/µl and 200 pg/µl. Even for 20 pg/µl samples the deviation is below 8.6%. For comparison, we calculated the concentrations of the DNA mixtures according to the uncorrected standard procedure. For this purpose, we used the 50 bp dilution series (see Fig 4) as calibration standard and calculated the concentrations of the DNA mixtures on the basis of the fitted calibration measuring points. Fig 7 shows a comparison of the corrected results versus the uncorrected results. The complete data of all mixtures can be found in S1 Table in the appendix. Without the bleaching correction, the actual concentration is significantly underestimated while the corrected calibration procedure provides much more accurate results.

**Fig 7.**
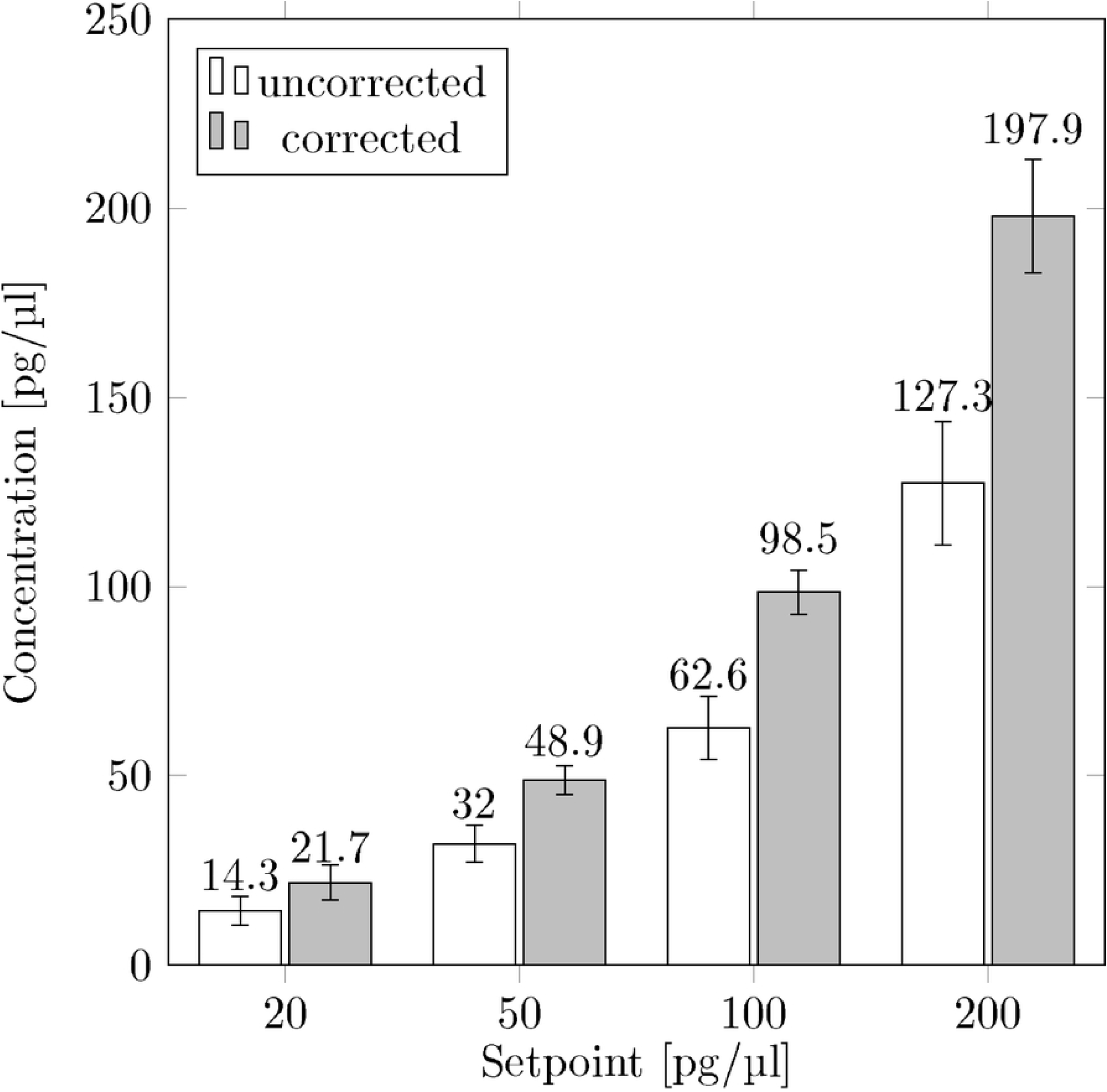
Results of the determination of mass concentration. The mass concentration of eleven DNA mixtures (2 µl drops) with setpoint concentrations of 20 pg/µl, 50 pg/µl, 100 pg/µl and 200 pg/µl are determined. Comparison of the new calibration procedure to uncorrected conventional procedure using directly fluorescence intensity of 50 bp dilution series (see figure 4) to determine the mass concentration.

We believe that our improved calibration method will make measuring molecular biological samples of unkown sequence composition effortless, accurate and sample-saving when compared to previous methods.

We would like to point out that besides photobleaching, the so-called saturation of optical intensity also plays an important role in fluorescence intensity. While the saturation of the fluorescence intensity only occurs at soaring laser intensities, significant parts of the molecules can already change into long-lasting triplet states at consideralby lower laser powers. Since our measurements were all taken at the same laser power, the relative deviation due to this effect is the same for all our measurements and can be neglected for correction.

## Conclusion

In this paper we derived a procedure to get highly accurate results for DNA measurements in highly diluted solutions. The challenge is to correct the so called photobleaching effect which reduces the fluorescence rate of the sample. The larger the hydrodynamic radius of the sample the larger is the photobleaching effect. In a first step, we determined the diffusion properties of the sample by means of fluorescence correlation spectroscopy. By doing this we could correct the fluorescence rate of the sample. In contrast to uncorrected measurements, we could reduce the photobleaching dependent failure of fluorescence based measurements to less than 3% compared to 30% without correction. Even for very low sample concentrations of 20 pg/µl the failure is still below 9%. This is remarkable considering that we conducted the measurements in tiny volumes of 2 µl. But even measurements in 1 µl droplet volumes are feasible. Which means that in case of 20 pg/µl our method only needs 20 pg of DNA to provide accurate results without the cost of expensive consumables. We would like to point out that for each mixture the average diffusion time is measured which enables the calculation of the average DNA fragment size of the sample. In principle, this allows a direct measurement of the sample molarity. It is also thinkable to determine the degree of fragmentation of nucleic acids in a sample. This opens up interesting fields of application in the field of DNA and RNA extraction from rare samples such as tissue sections. Furthermore, this circumvents time-consuming and expensive examinations of the sample using for example capillary electrophoretic methods.

## Supporting information

**S1 Table.** Data of mass determination. Mass determination of eleven mixtures with four defined mass concentrations (20 pg/µl, 50 pg/µl, 100 pg/µl and 200 pg/µl)

| Mixture | 20 pg/μl | 50 pg/μl | 100 pg/μl | 200 pg/μl |
| --- | --- | --- | --- | --- |
| 50 bp:1000 bp (1:1) | 21.01 | 47.77 | 111.38 | 214.21 |
| 50 bp:1000 bp (1:2) | 17.47 | 51.98 | 97.57 | 176.96 |
| 50 bp:1000 bp (1:3) | 20.51 | 49.29 | 92.91 | 210.87 |
| 50 bp:1000 bp (1:4) | 19.04 | 49.70 | 93.78 | 208.29 |
| 50 bp:1000 bp (1:5) | 19.62 | 44.80 | 96.71 | 190.54 |
| 50 bp:1000 bp (1:6) | 17.52 | 45.37 | 104.06 | 165.71 |
| 200 bp:500 bp (1:1) | 24.98 | 54.99 | 99.59 | 204.91 |
| 200 bp:500 bp (1:2) | 23.78 | 53.02 | 93.66 | 206.66 |
| 200 bp:500 bp (1:3) | 16.05 | 42.71 | 94.54 | 200.74 |
| 200 bp:500 bp (1:4) | 30.47 | 51.44 | 95.79 | 202.47 |
| 200 bp:500 bp (1:5) | 28.40 | 46.72 | 103.98 | 195.55 |
| Mean [pg/μl] | 21.71 | 48.89 | 98.54 | 197.90 |
| Variance coefficient [%] | 23.31 | 7.60 | 5.76 | 7.47 |
| Deviation from target[%] | 8.57 | -2.22 | -1.46 | -1.05 |

## Acknowledgments

We would like to thank all other involved members of the BioMOS group at Fraunhofer FIT for their support.

## References

1. Hoseini SS, Sauer MG. Molecular cloning using polymerase chain reaction, an educational guide for cellular engineering Molecular cloning using polymerase chain reaction, an educational guide for cellular engineering. Journal of Biological Engineering. 2015;9(2):1–12.

2. Robin JD, Ludlow AT, LaRanger R, Wright WE, Shay JW. Comparison of DNA Quantification Methods for Next Generation Sequencing. Scientific Reports. 2016;6(1):24067. doi:10.1038/srep24067.

3. Shi Sr, Cote RJ, Wu L, Liu C, Datar R, Shi Y, et al. DNA Extraction from Archival Formalin-fixed, Paraffin-embedded Tissue Sections Based on the Antigen Retrieval Principle: Heating Under the Influence of pH. The Journal of Histochemistry & Cytochemistry. 2002;50(8):1005–1011.

4. Doleshal M, Magotra AA, Choudhury B, Cannon BD, Labourier E, Szafranska AE. Evaluation and Validation of Total RNA Extraction Methods for MicroRNA Expression Analyses in Parafin-embedded Tissues. The Journal of Molecular Diagnostics. 2008;10(3):203–211. doi:10.2353/jmoldx.2008.070153.

5. Schmittgen TD, Livak KJ. Analyzing real-time PCR data by the comparative C T method. Nature Protocols. 2008;3(6):1101–1108. doi:10.1038/nprot.2008.73.

6. Lee C, Kim J, Shin SG, Hwang S. Absolute and relative QPCR quantification of plasmid copy number in Escherichia coli. Journal of Biotechnology. 2006;123:273–280. doi:10.1016/j.jbiotec.2005.11.014.

7. Epstein JR, Biran I, Walt DR. Fluorescence-based nucleic acid detection and microarrays. Analytica Chimica Acta. 2002;469(1):3–36. doi:10.1016/S0003-2670(02)00030-2.

8. Dragan AI, Casas-Finet JR, Bishop ES, Strouse RJ, Schenerman MA, Geddes CD. Characterization of PicoGreen interaction with dsDNA and the origin of its fluorescence enhancement upon binding. Biophysical Journal. 2010;99(9):3010–3019. doi:10.1016/j.bpj.2010.09.012.

9. Dragan AI, Pavlovic R, McGivney JB, Casas-Finet JR, Bishop ES, Strouse RJ, et al. SYBR Green I: Fluorescence properties and interaction with DNA. Journal of Fluorescence. 2012;22(4):1189–1199. doi:10.1007/s10895-012-1059-8.

10. Schwille P, Haustein E. Fluorescence correlation spectroscopy. An introduction to its concepts and applications. Fluorescence Correlation Spectroscopy. 2009; p. 1–33. doi:10.1002/lpor.200910041.

11. Song L, Hennink EJ, Young IT, Tanke HJ. Photobleaching kinetics of fluorescein in quantitative fluorescence microscopy. Biophysical Journal. 1995;68(6):2588–2600. doi:10.1016/S0006-3495(95)80442-X.

12. Enderlein J, Robbins DL, Ambrose WP, Goodwin PM, Keller RA. The statisticas of single molecule detection: An overview. Bioimaging. 1997;5:88–98.

13. Magde D, Elson E, Webb WW. Thermodynamic fluctuations in a reacting system measurement by fluorescence correlation spectroscopy. Physical Review Letters. 1972;29(11):705–708. doi:10.1103/PhysRevLett.29.705.

14. Petrov EP, Schwille P. State of the Art and Novel Trends in Fluorescence Correlation Spectroscopy. Standardization and Quality Assurance in Fluorescence Measurements II. 2008;6:145–197. doi:10.1007/4243_2008_032.

15. Rigler R, Mets Ü, Widengren J, Kask P. Fluorescence correlation spectroscopy with high count rate and low background: analysis of translational diffusion. European Biophysics Journal. 1993;22(3):169–175. doi:10.1007/BF00185777.

16. Aragón SR, Pecora R. Fluorescence correlation spectroscopy and Brownian rotational diffusion. Biopolymers. 1975;14(1):119–137. doi:10.1002/bip.1975.360140110.

17. Widengren J, Mets l, Rigler R. Fluorescence Correlation Spectroscopy of Triplet States in Solution: A Theoretical and Experimental Study. Journal of Physical Chemistry. 1995;99(36):13368–13379. doi:10.1021/j100036a009.

18. Ishii K, Tahara T. Correction of the afterpulsing effect in fluorescence correlation spectroscopy using time symmetry analysis. Optics Express. 2015;23(25):32387. doi:10.1364/OE.23.032387.

19. Petrašek Z, Schwille P. Precise Measurement of Diffusion Coefficients using Scanning Fluorescence Correlation Spectroscopy. Biophysical Journal. 2008;94(4):1437–1448. doi:10.1529/biophysj.107.108811.

20. Doi M, Edwards SF. The theory of polymer dynamics. vol. 73 of International series of monographs on physics. Reprint ed. Oxford: Clarendon Press; 1986.

21. Nkodo AE, Garnier JM, Tinland B, Ren H, Desruisseaux C, McCormick LC, et al. Diffusion coefficientof DNA molecules during free solution electrophoresis. Electrophoresis. 2001;22:2424–2432.

22. Robertson RM, Laib S, Smith DE. Diffusion of isolated DNA molecules: Dependence on length and topology. Proceedings of the National Academy of Sciences. 2006;103(19):7310–7314. doi:10.1073/pnas.0601903103.

23. Zander C, Enderlein J, Keller R. Single molecule detection in solution: Methods and applications. 1st ed. Zander C, Enderlein J, Keller RA, editors. Berlin: Wiley-VCH; 2002. Available from: http://dx.doi.org/10.1002/3527600809.

